# A Comparison of Lossless Compression Methods in Microscopy Data Storage Applications

**DOI:** 10.1101/2023.01.24.525380

**Authors:** Logan A. Walker, Ye Li, Maggie McGlothlin, Dawen Cai

**Affiliations:** University of Michigan, Ann Arbor, Michigan, USA

**Keywords:** microscopy, data management, compression

## Abstract

Modern high-throughput microscopy methods such as light-sheet imaging and electron microscopy are capable of producing petabytes of data inside of a single experiment. Storage of these large images, however, is challenging because of the difficulty of moving, storing, and analyzing such vast amounts of data, which is often collected at very high data rates (>1GBps). In this report, we provide a comparison of the performance of several compression algorithms using a collection of published and unpublished datasets including confocal, fMOST, and pathology images. We also use simulated data to demonstrate the efficiency of each algorithm as image content or entropy increases. As a result of this work, we recommend the use of the `BLOSC` algorithm combined with `ZSTD` for various microscopy applications, as it produces the best compression ratio over a collection of conditions.

**CCS CONCEPTS:** • Applied computing → Bioinformatics; Imaging.

## 1 INTRODUCTION

The biomedical sciences have been fundamentally altered by the introduction of ever-improving imaging techniques, such as confocal [26, 27], light-sheet [3, 10, 21, 24], super-resolution microscopy [11, 18, 23], and expansion microscopy [20, 22]. As microscopes become faster and higher resolution, their utility have increased. Some of the applications of these methods in neuroscience include multi-spectral imaging of Brainbow samples to identify neuronal connections [5, 20], fMOST imaging for whole-brain neuron reconstruction [4, 17], and single molecule imaging techniques for spatial transcriptomics, which allow scientists to identify spatial profiles of RNA expression [14]. As experiments become more content-rich and data sizes increase, it is increasingly difficult to handle the data, which may can contain many single images reaching >10TB [9, 26]. To deal with data of this scale, most microscopy images are stored in some form of compression, however, there is little consensus on the best way to perform compression to optimize both the compression ratio and the data read/write speed. For archival and distribution, it is not uncommon to see data distributed as compressed TIFF files which have been processed using `GZIP` or `ZIP`, because these tools are commonly-available on most desktop computer and integrated with various image visualization tools. In other cases, matrix storage libraries like Zarr [15] or HDF5 [8, 12] are used to store images in a chunked format which can then be compressed using a variety of methods. Despite this variability, there have been few efforts to benchmark the best compression practices in microscopy data storage. In one recent example, Datta and colleagues showed that different compression algorithms for storage of reduced-representation electron microscopy can have significantly different performance [7], showing the importance of this type of comparison for designing experimental protocols. Here, we will perform benchmarking of many of the most common lossless image compression algorithms, including `GZIP`, `LZ4`, `BLOSC`, and `ZSTD` using a variety of real-world and simulated biomedical images. From these results we provide a recommendation for compression of biomedical images.

## 2 DATASETS

### 2.1 Imaging data

We collected 9 different datasets from a variety of sources and imaging modalities for benchmarking, including published and unpublished data (Table 1). As specified in the table, each dataset were assigned a label to make their referencing in the text simpler and the collection of datasets spans many orders of magnitude in size. The published datasets include traditional confocal imaging of Brainbow-labeled neurological tissue (`brainbow`) [5, 17], fMOST imaging of an entire mouse brain (`fMOST`) [26], and breast cancer tissue pathology images collected as part of a 10 `×`visium experiment (`visium`). In addition to this, we collected confocal imaging of DiDdyed neurites from mouse cortical tissue (`neurites`), 5-channel *Drosophila melanogaster* (fruit fly) Bitbow images as previously described in [13] (`bitbow`), and fluorescent latex beads (`bead`). We also include an example segmentations from the mouse common coordinate framework (CCF; `segment`) [25], which is a categorical image representing the locations of different brain regions; this is a meaningful comparison because categorical segmentation images are commonly used to identify structures in different micrographs (e.g., [6]). Negative control images of background noise collected from an ORCA-Fusion sCMOS camera (Hammatsu Photonics K.K.) are included at both 8 and 16 bit (`noise8`,`16`), because they represent the lowest-content image that can be produced by a typical microscopy system. Figure 1 displays example frames from each image.

**Table 1:** A list of the datasets collected for this study, with associated metadata. Image size is given in *c* × *z* × *x* × *y*.

| Label | Size (px) | Type | Source |
| --- | --- | --- | --- |
| brainbow | $4 \times 136 \times 512 \times 512$ | uint16 | [17] |
| neurites | $1 \times 1000 \times 2304 \times 2304$ | uint16 | (this work) |
| bitbow | $5 \times 97 \times 3000 \times 3000$ | uint16 | (this work) |
| fMOST | $2 \times 11464 \times 20821 \times 30801$ | uint16 | [16] |
| visium | $3 \times 1 \times 24240 \times 24240$ | uint8 | 10xgenomics.com |
| bead | $1 \times 1000 \times 2304 \times 2304$ | uint16 | (this work) |
| noise16 | $1 \times 100 \times 1000 \times 1000$ | uint16 | (this work) |
| noise8 | $1 \times 100 \times 1000 \times 1000$ | uint8 | (this work) |
| segment | $1 \times 456 \times 528 \times 320$ | uint32 | [25] |

**Figure 1:**
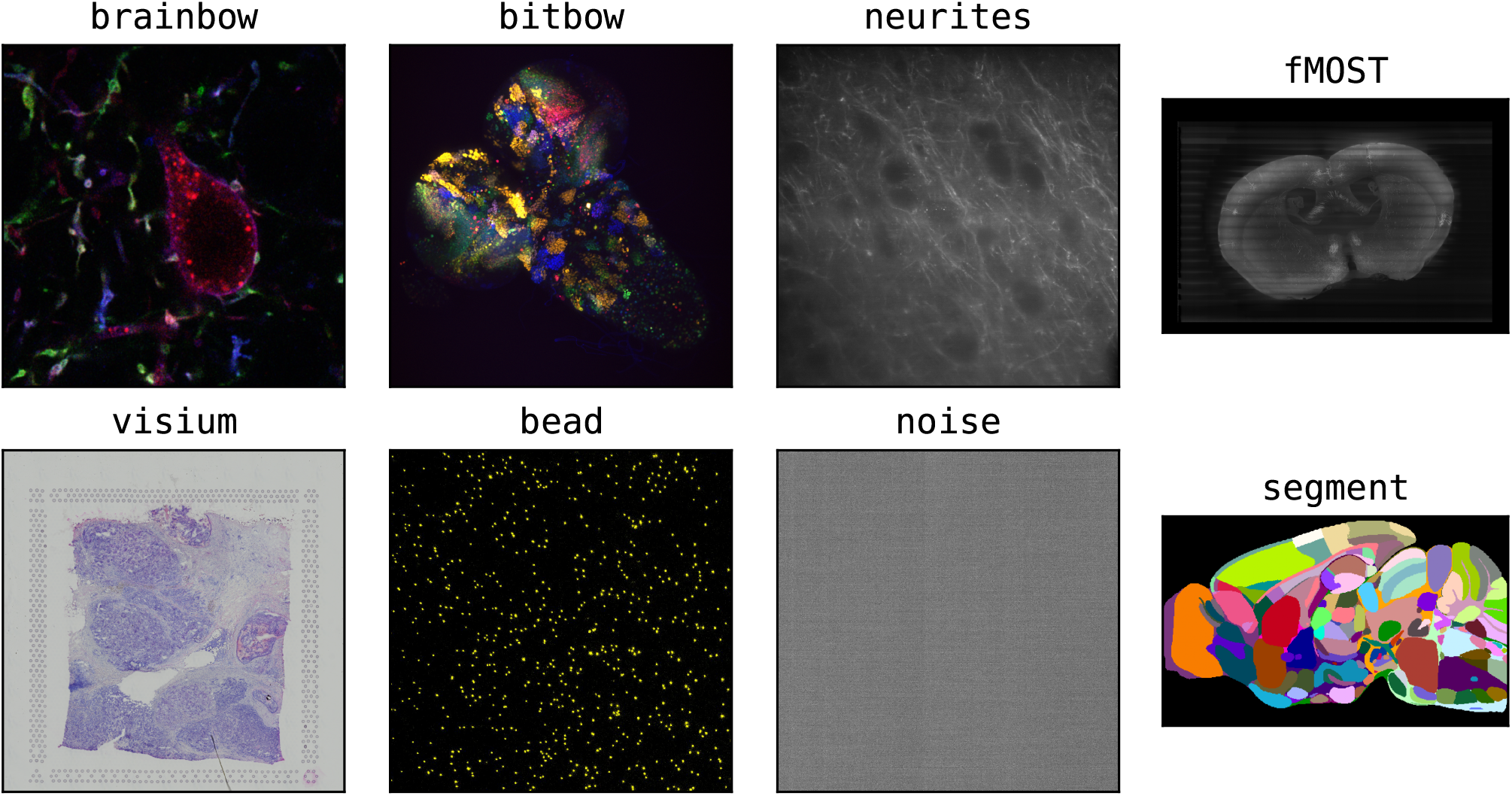
Example z-axis slices from each dataset presented in Table 1. In some images, maximum projections have been performed to increase the visibility of the image in print.

### 2.2 Simulated Data

In addition to the real datasets presented above, we used custom Python code to generate *in silico* confocal images of simulated point sources (beads), randomly distributed throughout the frame(see **Methods**). In total, we created 27 images of (256 px) 3 at 9 logarithmically-spaced bead densities which were used for titration experiments at a signal-to-noise-ratio (SNR) of 5.

## 3 METHODS

### 3.1 Compute Environment

Dataset compression tests (**Section 4.1**) were performed on a custom-built server with two EPYC 7351 16 core processors and 512GB of DDR4 memory running at 2133 MT/s. This system was chosen because it is representative of real-world situations, where high performance computing is needed to handle data of this scale. Simulation results (**Section 4.2**) were performed on a MacBook Pro with a M1 Pro processor and 16GB of LPDDR5 memory at 6400 MT/s, because of it’s higher speed than the sever described above. We forced all benchmarks to run on a single core, because some compression methods are unable to be multithreaded, but we note that, in practice, many datasets can be compressed in multiple streams to achieve higher throughput than we present here. All files were saved into a Python memory buffers rather than a hard drive in order to ensure that disk latency or bandwidth did not impact benchmark times. As such, we use a parameter set labeled “Uncompressed” as a control for the throughput of the testing scripts themselves. We used Anaconda Python (v3.10) to implement our testing scripts, as it provides a high-level programming interface, while having a minimal impact on performance because the libraries used in this study are implemented as wrappers around the low-level compression libraries.

Compression methods tested included `LZ4`, `LZF`, `GZIP`, `ZSTD`, and `BLOSC` [1]. For methods which accept compression level parameters, we tested a range of parameters between the minimum and maximum. BLOSC uses a “meta-compression” and blocking strategy which allows the use of multiple compressors as well as filters, such as `bitshuffle` and `shuffle`. These images transpose the data at the bit or byte level in order to align similarly-value bits in integer data, and require a negligible amount of computation to perform. We excluded comparisons without `BLOSC` shuffling filters, because we found that they consistently performed worse than the shuffled equivalent (data not shown). In the rest of this report, we will describe each compression method using the combination of parameters used for each comparison.

### 3.2 Simulated data generation

We used custom Python scripts to generate simulated confocal images of varying complexity using the `python-sdt` simulation package [19]. Specifically, random 3D bead coordinates were generated for each specified density, which were then modeled as point sources with an intensity of 500 au and standard deviations of 2 px. Each 2D frame’s coordinates were then calculated by slicing the 3D bead coordinates into individual planes assuming a Gaussian intensity distribution for each point. The image was then offset by 100 au and noise was added by sampling with the Poisson distribution with λ equal to the simulated image intensity:

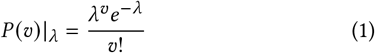

Each 2D frame is then stacked to form composite 3D images which were exported for benchmarking.

### 3.3 Compression comparison

Each dataset was sampled into 10 independent 256 ×256 ×256 px chunks across all channels in *x*× *y* ×*z*. Each sampled image was compressed into HDF5 files with a chunk size of 32× 32× 32 px using default parameters, with the exception of the compression settings which were changed for each experiment. We quantify compression and decompression rates as an “Effective” throughput, which is the size of uncompressed data divided by the total time to open, parse, and decompress an image into RAM. This allows us to quantify the speed that a given application will be able to actually access data, rather than just the algorithm speed. The chunk size was reduced for the `visium` dataset, because the image is a single z plane. We used compression filters for HDF5 which were included in the base Python HDF5 (`h5py`1) library or the `hdf5plugin`2 `pip` package, which extends the number of available compression plugins available.

## 4 RESULTS AND DISCUSSION

### 4.1 Compression Benchmark

The compression ratios (c.r.), decompression, and compression speed of each compression experiment are shown in Figure 2. For visualization, we removed the `segment` layer because it had very high compression rates which were not representative of the other datasets (c.r. range 11.3-96.6) with `ZSTD`-based approaches performing the best. First, we observe `noise16` consistently returns the largest compression ratio, while `noise8` performs poorly due to it’s much higher relative entropy. `visium` also performs poorly, likely due to the 2D nature of the dataset causing smaller chunking sizes. In this sorted figure, we see that the shuffle and bitshuffle filters are essential for achieving the highest compression ratios in integer data. `BLOSC-ZSTD-SHUFFLE-8` is found to be the highest compression ratio by a very small margin, but we recommend the use of `BLOSC-ZSTD-SHUFFLE-5` when high compression ratios are needed because the higher compression speed found is much higher.

**Figure 2:**
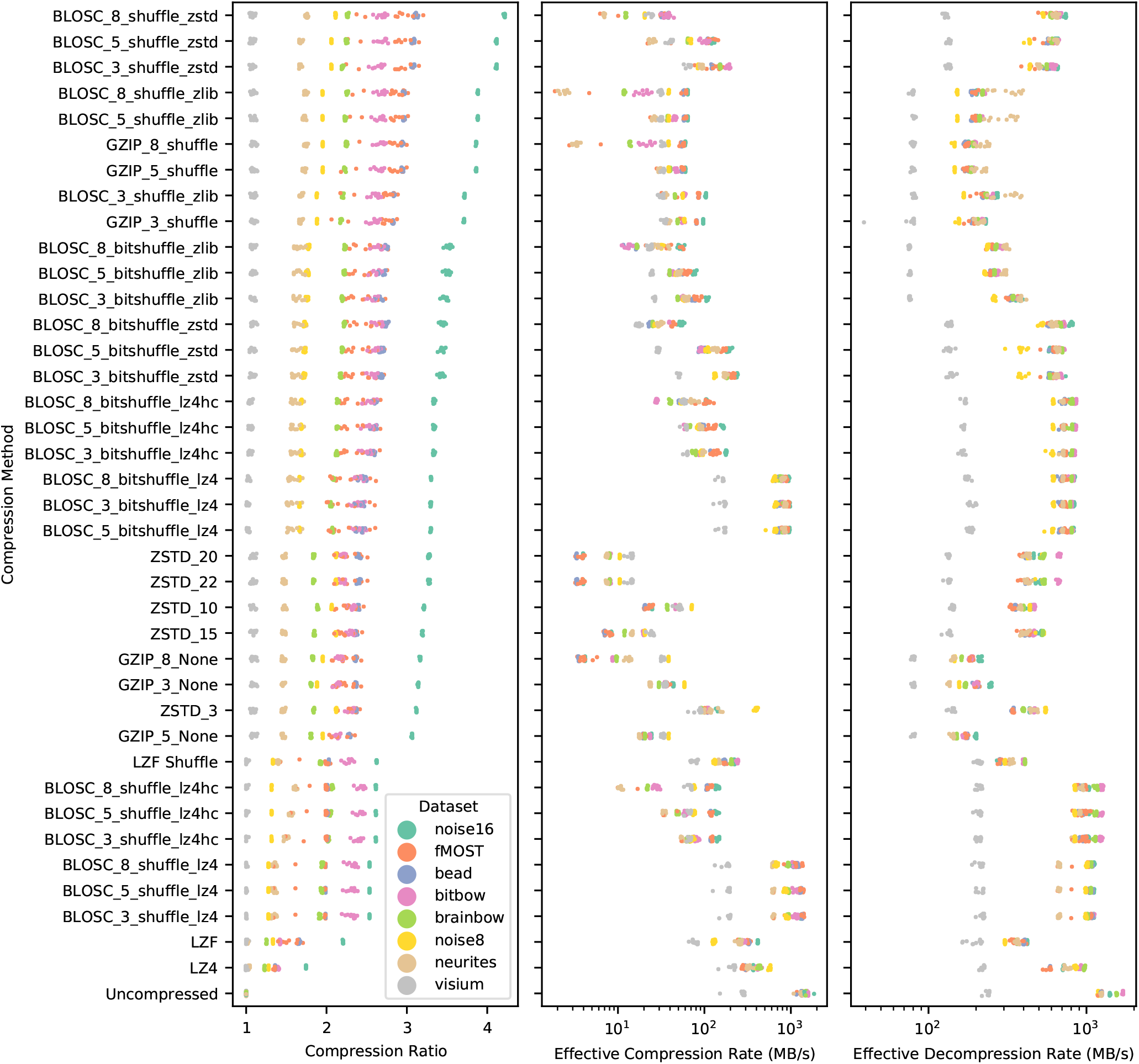
A strip plot of the compression ratio achieved by different compression algorithms when applied to our benchmark datasets. Methods are sorted by the maximum compression rate achieved by any dataset for visualization.

Next, we compared the compression and decompression speed of a subset of compression methods which were selected to represent the full dynamic range of speed and compression ratios (Figure 3). Here, we can see that the algorithms using `LZ4` are universally the fastest methods, with `LZ4` and `BLOSC-LZ4-BITSHUFFLE-5` performing only slightly slower than a the uncompressed control. We highlight these methods as a good candidates for real-time data processing, such as intermediate compression of data being produced by a microscope due to these results, with the `BLOSC` variant being prefered due to the higher compression ratio (Figure 2). The `BLOSC-ZSTD-SHUFFLE-5` method identified above is able to be decompressed quickly, however it’s compression rates are slow enough to be a limiting for real-time applications. `GZIP` is too slow to be usable by in any real-time microscopy application.

**Figure 3:**
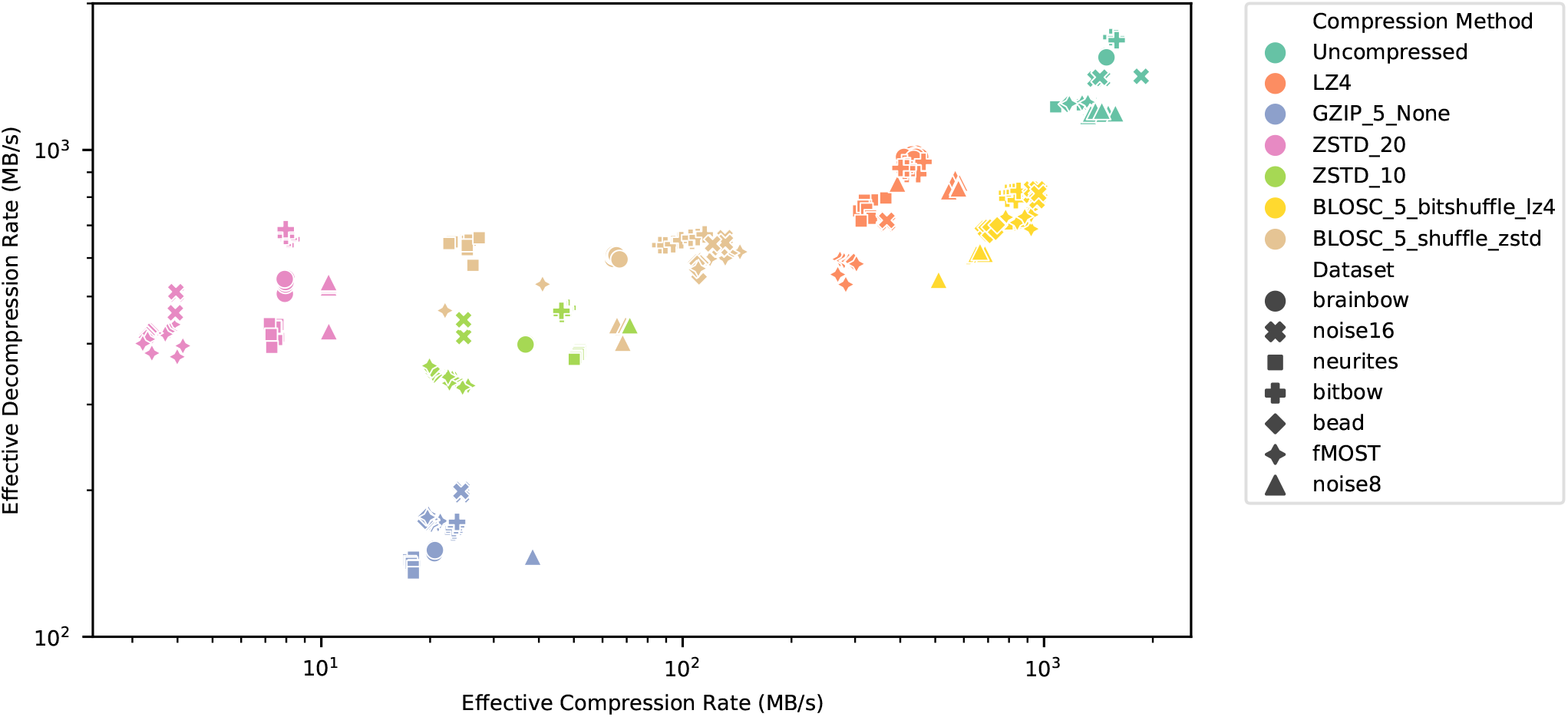
A scatter plot of the compression compression and decompression rates for the different compression algorithms tested.

### 4.2 Simulated Confocal Imaging Benchmark

We created simulated micrographs with varying densities of fluorescent beads, similar to the `beads` dataset and with simulated image noise (Figure 4, left). We compared the compression ratios of `BLOSC-ZSTD-BITSHUFFLE-5` with a commonly-used standard compression scheme represented by `GZIP-5`. Here, we find that `BLOSC` performs significantly better, however there is a reduction in the compression ratio as the bead count increases. This matches our expectation, as increased bead density is directly correlated with an increase in image entropy, making the theoretical compression limit lower.

**Figure 4:**
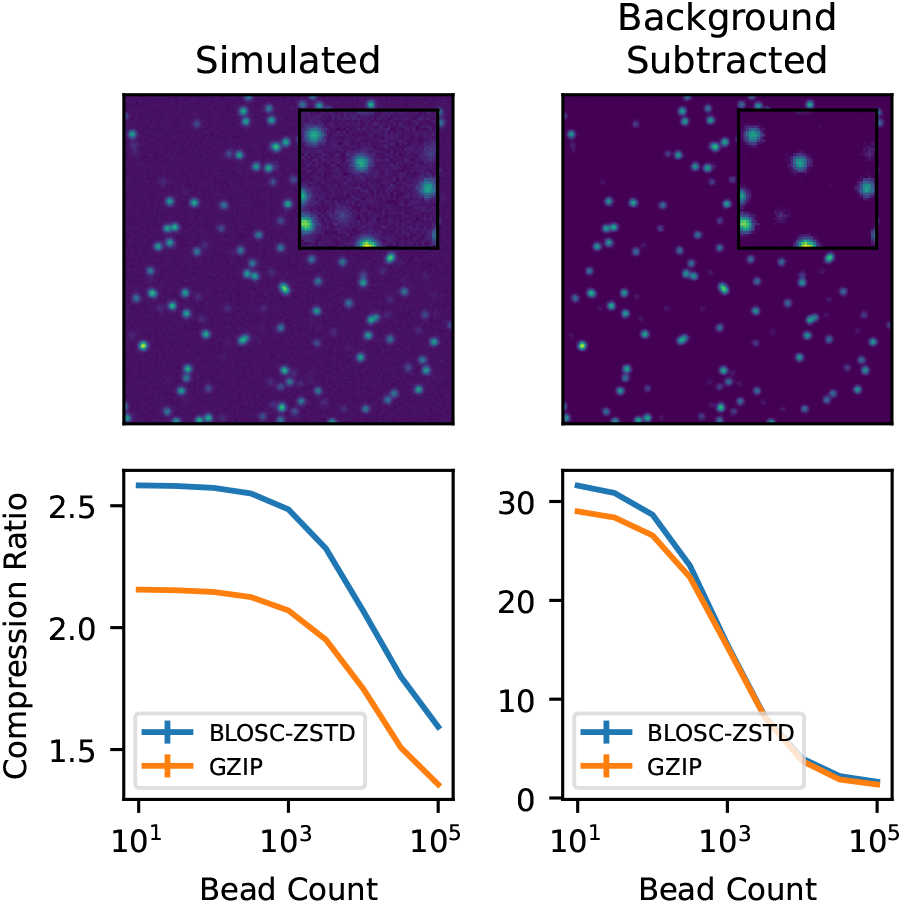
Performance of BLOSC-ZSTD and GZIP compression at compressing simulated confocal data.

Next, to demonstrate the effectiveness of simple data manipulation for improving we performed background subtraction above the approximate noise floor of the images described above (120au; Figure 4, right). We repeated the compression testing, finding that there is an order-of-magnitude increase in the achieved compression ratio, with `GZIP` performing better than `BLOSC` in this case. While this transform is lossy, it highlights the potential value of preprocessing microscopy data before storage, as well as the potential for other lossy compression methods.

## 5 CONCLUSION AND FUTURE DIRECTION

Compression is incredibly important for making biomedical imaging both economically and, in many cases, technically feasible by reducing the hard disk space needed for a given experiment. We have shown benchmarks of 39 different compression parameter combinations applied to 9 different biomedical image datasets derived from different neurological tissues and imaging methods, as well as simulated data. Through this quantification, excluding `segment` and `visium` due to their outlier characteristics, we find that `BLOSC-ZSTD-SHUFFLE-5` performs the best of all methods compared (c.r. 2.7± 0.8), however, at the expense of slow compression (86 ±36 MB/s). In cases where fast or real-time computation is needed, `BLOSC-LZ4-BITSHUFFLE-8` provides a good compromise (c.r. 2.3 ±0.5) while being fast to compress (790 ±92 MB/s). These results provide an important starting point for choosing a compression method for any specific experiment, and we provide our code under an GPL license to allow testing on different imaging types or CPU architectures, as we expect results may vary.While we presented a comprehensive view of the applications of lossless compression in microscopy, there are several factors which should be considered as future directions or factors when deciding which compression method to use for a specific microscopy application. First, NVIDIA has recently developed toolkits for lossless compression which take advantage of GPU compression, potentially making much higher throughputs possible by application of these tools^3^. Next, while we used a chunk size of (32px) ^3^ in this study because it allows quick access in all dimensions, it is possible that different datasets may have different spatial correlations which make increased chunk sizes more efficient. Finally, as shown in Section 4.2, image filtering and preprocessing has a strong potential for improving the efficiency of data compression, assuming you can make guarantees about the scientific validity of the decompressed data. In the future, various lossy compression methods may be able to fill this gap, similar to the previously-reported B^3^D method [2].

## DATA AND CODE AVAILABILITY

The code used in this article is available from the Cai Lab GitHub page under a GPL license^4^. The novel images presented in this comparison have been uploaded to the University of Michigan Deep Blue Data repository (DOI pending) and are freely available under a Creative Commons Attribution-ShareAlike 4.0 International (CC BY-SA 4.0) license^5^.

## AUTHOR CONTRIBUTIONS

LAW and DC conceived this project. LAW wrote associated analysis code and first draft of the manuscript. LAW, YL, and MM performed biological sample preparation and microscopy for novel datasets used in analysis. All authors edited and approved the final manuscript.

## ACKNOWLEDGMENTS

LAW was supported by a University of Michigan Rackham Predoctoral Fellowship. This work was supported by NIH-RF1MH123402, and NIH-RF1MH124611 to DC.

## Footnotes

1 https://www.h5py.org/

2 https://pypi.org/project/hdf5plugin/

3 https://github.com/NVIDIA/nvcomp

4 https://github.com/Cai-Lab-at-University-of-Michigan

5 https://creativecommons.org/licenses/by-sa/4.0/

